# Mechanical Properties of a Primary Cilium from the Stochastic Motions of the Cilium Tip

**DOI:** 10.1101/292409

**Authors:** J. Flaherty, Z. Feng, Z. Peng, Y.-N. Young, A. Resnick

## Abstract

The stochastic tip dynamics of a primary cilium held within an optical trap is quantified by combining experimental, analytical and computational tools. Primary cilia are cellular organelles, present on most vertebrate cells, hypothesized to function as a fluid flow sensor. The mechanical properties of a cilium remain incompletely characterized. We measured the fluctuating position of an optically trapped cilium tip under untreated, Taxol-treated, and HIF-stabilized conditions. We applied analytical modeling to derive the mean-squared displacement of the trapped tip of a cilium and compared the results with experimental measurements. We provide, for the first time, evidence that the effective flexural rigidity of a ciliary axoneme is length-dependent, and longer cilia are stiffer than shorter cilia. We then provide a rational explanation for both effects. We demonstrate that the apparent length-dependent flexural rigidity can be understood by a combination of modeling axonemal microtubules orthotropic elastic shells and including (actin-driven) active stochastic basal body motion. It is hoped that our improved characterization of cilia will result in deeper understanding of the biological function of cellular flow sensing by this organelle. Our model could be profitably applied to motile cilia and our results also demonstrate the possibility of using easily observable ciliary dynamics to probe interior cytoskeletal dynamics.

## INTRODUCTION

Primary cilia are slender hair-like structures, several microns long and 0.2 microns in diameter, present on most vertebrate cells, that protrude from the cell body into the extracellular space. Long considered vestigial structures, recent work has conclusively demonstrated that the primary cilium is in fact a signaling center for the cell (1) organizing a large number of signaling pathways. Demonstrations that bending a primary cilium via fluid flow (2), optical tweezers (3), or micropipette (4) initiates intracellular calcium release imply that physiologically, the primary cilium is a flow sensor. However, the biological significance of this function remains unclear in part due to incomplete understanding of the dynamics of the cilium in the presence of flow (5–7). For example, a recent report (7), called into question the proposed mechanism by which flow (kinetic) energy is transduced into biochemical (potential) energy. Because bending the cilium involves mechanical stress and strain, one of our research foci centers on characterizing the mechanical properties of the primary cilium and exploring ways to modify the flow response by pharmacologically altering the mechanical properties of the primary cilium.

The primary cilium ultrastructure has been well characterized (8), and most mechanosensation work has focused on ciliary associated transmembrane proteins, for example polycystin 1, polycystin 2 (9) and polycystin related proteins PKD1L1 and PKD2L1 (1). By contrast, the possible role of structural elements in ciliary mechanotransduction has received less attention. Primary cilium structure consists of 9 microtubule doublets anchored in the basal body, which itself is a highly organized structure comprising a centrosome (microtubule triplets), transition fibers, a rootlet, and the basal foot (10). We hypothesize that the mechanosensing function of cilia can occur by straining these microtubule structural elements in a similar manner to actin-mediated mechanosensation (11). Indeed, preliminary results indicate that the basal body may have a role in differentiating mechanosensation from chemosensation (2, 12, 13).

### Experimental Rationale

Here, we consider the primary cilium as a passive mechanical beam capable of transducing kinetic energy of flowing fluid into elastic strain (bending) energy. Bending the cilium seems to be required to initiate calcium signaling (2), and the amount of bending in response to applied flow can be characterized by the flexural rigidity (equivalently, bending modulus) (14) ‘EI’. Even so, the mechanical response of primary cilia to external loading remains incompletely characterized (15). Most prior measurements of the ciliary bending modulus fall within the range of 10-50 pN* *μ*m^2^ (14), (16), but there are also measurements that report significantly higher bending modulus, as high as 300 pN* *μ*m^2^ (17). As we will show and discuss, our results potentially resolve this discrepancy but also raise important new questions. It is essential to note that prior to this report, all reported measurements of the ciliary bending modulus, with the excepton of (15), calculate the bending modulus based on fitting an image of a bent cilium to the equation for a homogeneous cantilevered beam (discussed below, in ‘Analysis’). Our approach is qualitatively different and does not rely on imaging the cilium.

Measuring the strained position of the cilium tip provides information about the mechanical response of the cilium to an applied mechanical load. For example, subjecting a uniform cantilever of length ‘L’ to a static load ‘q’ localized to the free end results in a static tip displacement y = q L^3^/3EI. When both L and y are known, the expression can be inverted to calculate an ‘effective’ bending modulus EI_*eff*_ for example, EI_*eff*_ = q L^3^/3y.

Now, consider a set of cantilevered beams with varying ‘L’, each subjected to identical localized tip loads. If the tip displacement y does not scale as L^3^ but rather some other function of cantilever length (say L^*α*^), the cantilevered beam would be characterized as having a length-dependent bending modulus (see, for example, (18)). If the cantilever has a uniform cross-section, the ascribed length-dependence may simply reflect more complicated mechanics of the beam.

Consequently, we calculated EI_*eff*_ as a function of cilium length L and determined that the cilium has an apparent length-dependent bending modulus. Furthermore, we determined EI_*eff*_ when either Taxol, a microtubule stabilizer, or the small molecule CAY10585 is added to the culture medium. Hypoxia-inducible factor HIF-1 is destabilized by CAY10585 (19) and preliminary results demonstrate a link between ischemia-hypoxia and cilium bending modulus (20).

After presenting the data, we provide several attempts to determine the source of this apparent length dependence by: 1) Use of different boundary conditions; 2) Careful accounting of viscous forces along the axoneme; 3) Allowing the bending modulus to vary with position; 4) allowing the basal body to undergo active fluctuation; and 5) modeling the cilium as a collection of elastic shells rather than a homogeneous beam. We will show that a combination of improved structural modeling (orthotropic shell model) and stochastic modeling of ATP-driven actin cortex fluctuations of the basal body (21) reconcile our experimental data. We propose that this report provides novel insight into the mechanical response of the primary cilium to applied fluid flow, the notion of flow sensitivity by ciliated cells, and possibly the mechanism by which motile cilia generate fluid flow.

## MATERIALS AND METHODS

### Experimental Setup

Our experimental protocols for the growth, maintainance, and pharmacological manipulations of ciliated epithelial cells and optical trapping of cilia have been published elsewhere (15, 20, 22), so here we only provide a brief summary. Conditionally immortalized epithelial cells originally microdissected from the cortical collecting duct of an Immortomouse are grown to confluence and allowed to differentiate for several days, during which time a cilium emerges. A cultured epithelial tissue monolayer is then placed within an upright microscope sample incubation chamber for imaging and trapping. Trapping is carried out with the cells held at 39°C. The trapping beam enters the microscope from a side port, and the forward scattered trapping light is detected by a quadrant photodiode (QPD). The wavelength of the trapping beam (1064 nm) is well-tolerated by the cells (23, 24). The microscope objective used for the experiments was a 63X NA 0.90 dipping objective designed to be directly immersed into the culture media. The trapping geometry is known as a single-beam gradient trap, as a single beam is sufficient to confine the trapped object in all three dimensions. For our trap the beam waist is 0.3 *μm* and Rayleigh length is 0.4 *μ*m.

A schematic and representative image of an optically trapped primary cilium is shown in Figure 1. The cilium projects above the cell body, leaving the cells out of focus. As seen in the inset figure, the cilium appears as an in-focus dot against a blurred background.

**Figure 1:**
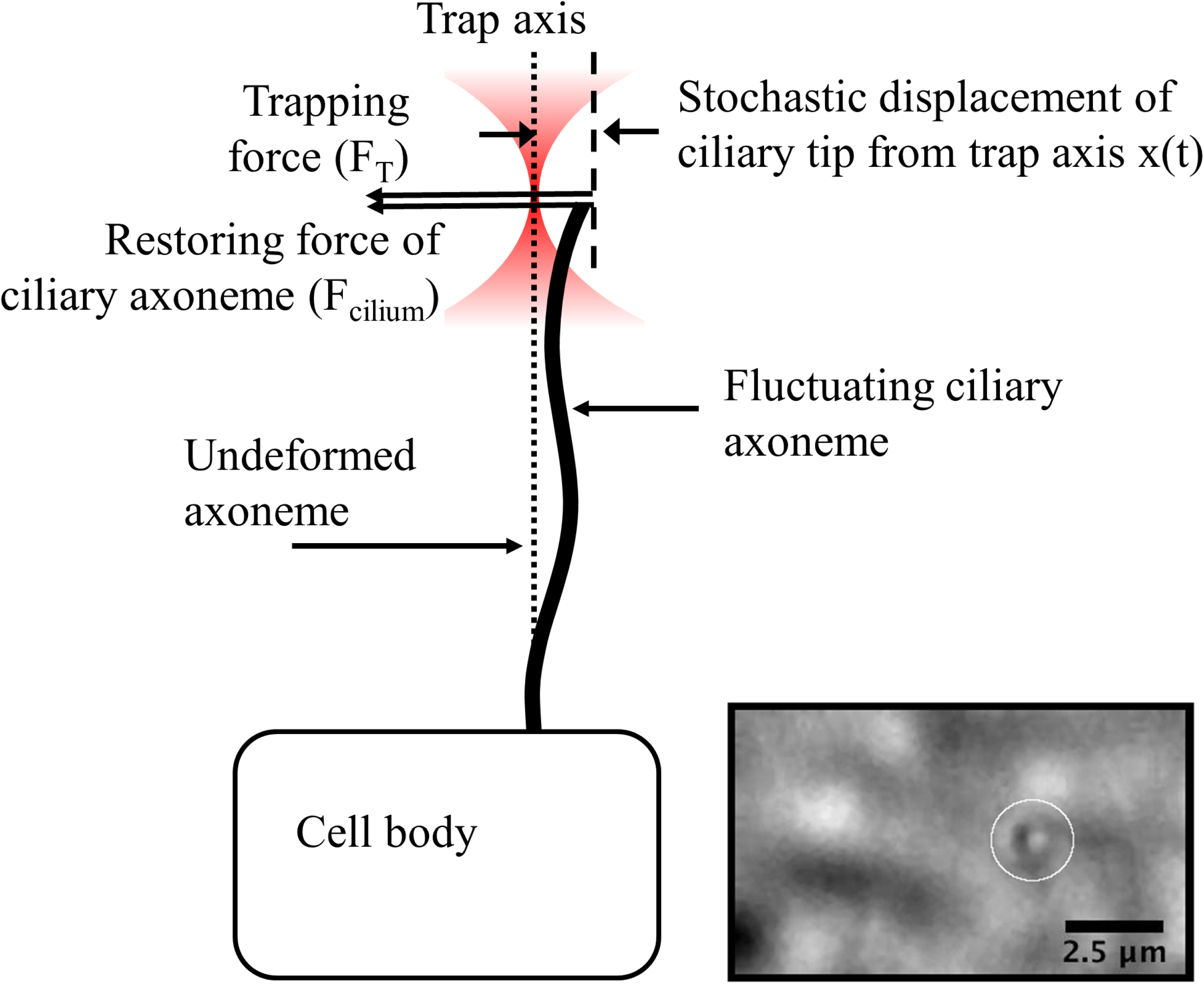
Schematic and inset image of optically trapped primary cilium. Location of optical trap indicated by circle. The cilium projects above the cell body and so appears as a dot in the microscopic image.

Experiments proceeded as follows: with the trapping beam turned off, a cilium is located and the length measured optically, relying on the thin depth of focus of the microscope objective and accurate z-stage positioning. The distal tip is positioned at the trap location, the trap turned on and QPD output sampled for several tens of seconds (sample rate = 10^4^ Hz). The tip of the cilium scatters the trapping beam, and as the tip moves stochastically due to Brownian motion, the angular distribution of scattered light changes. QPD data consists of a time-series of voltage samples, say 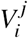, where i = 1,2,3,4 (each quadrant) and j is the sample number. For each sample, the 4 quadrants are combined to determine the location of the centroid along the ‘x’ and ‘y’ axis (axes are set by the rotational orientation of the QPD and fixed, but arbitrary with respect to the x- and y-axes of the camera), and the dataset 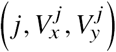 is considered ‘raw data’. The time-series raw data is then processed to calculate the mean-squared displacement (MSD) of the distal tip. As we will show, cilium length and asymptotic value of MSD are sufficient to characterize the effective bending modulus of a cilium in terms of an effective spring constant.

A good overview of various optical trap analysis methods can be found in (25). Our analysis method computes the MSD and fits an analytic function (presented below) to determine the trap stiffness. Analysis based on the asymptotic value of MSD is preferred for our system for two reasons. First, discretization of the QPD signal does not generate spurious results (26) and second, precise knowledge of viscous damping, which is problematic as our trapped object is a slender cylinder and not a sphere, is not required.

### Computational Model

We also developed a coarse-grained computational model of the primary cilium to study the deformation and fluctuations of the cilium axoneme and the basal body based on dissipative particle dynamics (DPD) (27). DPD is a coarse-grained molecular dynamics which aims to capture thermal fluctuations and hydrodynamic behaviors of the original atomistic systems (28, 29). Besides *non-bonded* DPD interactions to capture fluctuations and hydrodynamics, we apply coarse-graining to construct *bonded* interactions within the molecular structures of the primary cilium based on the state-of-the-art understanding of its structure. The details of the model is described in the SI text.

### Calculation of trapped object MSD

Data analysis is done using a Matlab procedure *QPD analysis Bulk* (22) that uses time-series data collected by the Quadrant Photodiode (QPD), which measures the x- and y-positions of the centroid of light scattered by a trapped particle. The algorithm calculates the autocorrelation function 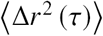 from a trajectory r(t) discretized into N_*T*_ + 1 time steps consisting of N_*A*_ = N_*T*_ - *τ* + 1 overlapping time intervals of duration *τ*:

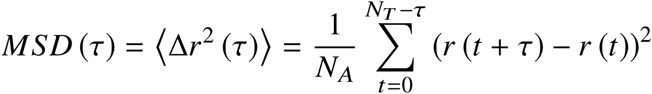

By fitting MSD(*τ*) to a known function (see subsections covering stochastic models) we obtained the long-time asymptotic limit value, which we denote MSD_∞_.

## RESULTS AND DISCUSSION

We present our data first and then follow with model-based interpretation of the data.

### Experimental measurements

#### Length-dependent bending modulus

We present our calculated values of the spring constant as a function of cilium length in Figure 2. The spring constant is related to the (effective) bending modulus of a trapped cilium, and the resulting calculated values of effective bending modulus are also presented in Figure 2.

**Figure 2:**
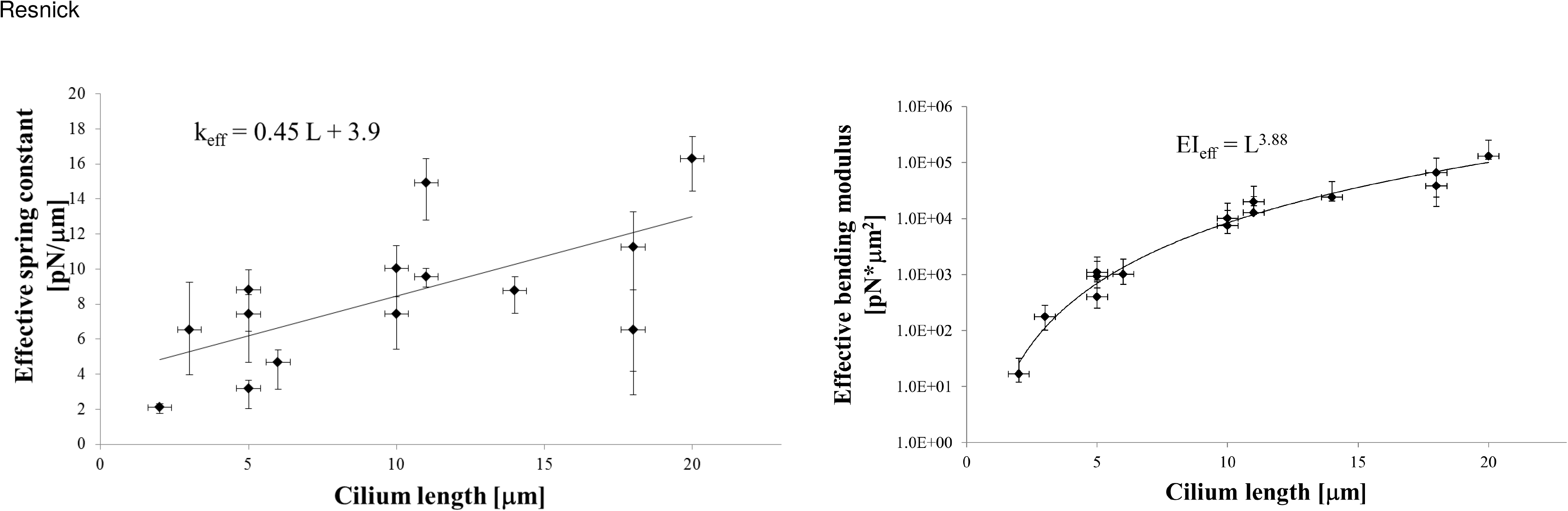
Calculated values of the trap spring constant and effective bending modulus as a function of cilium length

#### Pharmacological manipulation of cilia

We have previously investigated pharmacological alterations of the ciliary bending modulus (20) and here we present new data (Figure 3) showing similar effects. We examined the effects of two drugs, Taxol (paclitaxel) and CAY10585. Taxol has well understood effects on the mechanical stiffness of microtubules (30). CAY10585 is a small molecule that destabilizes Hypoxia-Inducible Factor-1 (HIF-1), a transcriptional regulator of hypoxia responses. CAY10585 also inhibits HIF gene transcriptional activity. We discuss the two treatments separately:

**Figure 3:**
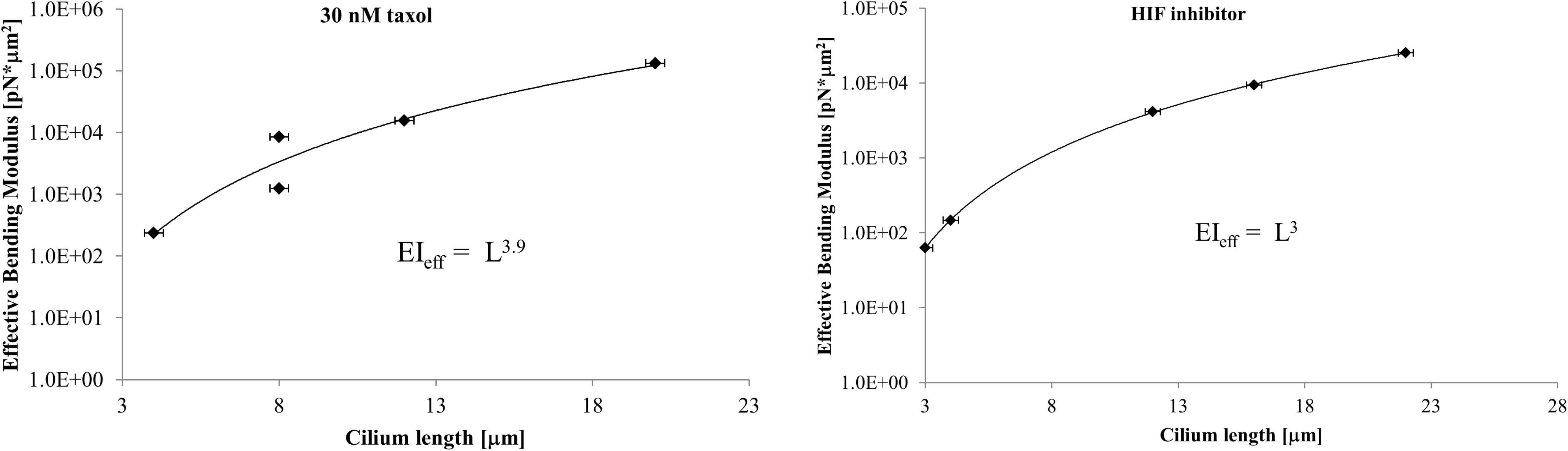
Calculated bending modulus values as a function of cilium length

#### Taxol

Taxol is a microtubule stabilizer, yet prior studies applying Taxol to microtubules (31) and cilia (20) show that the flexural rigidity decreases, a counterintuitive result. Based on molecular dynamics simulations, it was proposed (32) that Taxol increases axonemal stability by allowing the increased flexibility to relieve internal stresses. Figure 3 presents our data obtained by adding 30 nM of Taxol to the culture medium for 24 hours. Note that our applied concentration is well below earlier studies.

#### Hypoxia-Inducible Factor (HIF) inhibitor

We and others have previously examined the connection between localized hypoxia and primary cilium function. Hypoxia-Inducible Factors (HIFs) are potent transcription factors that control cellular responses to decreases in available oxygen. Both ciliary length and bending modulus have been shown to be altered when HIF is stabilized (20, 33). While the precise mechanism is currently unknown, the implications for altered flow sensitivity are profound; hypoxia may inhibit ciliary flow sensing at the initial transduction event rather than inhibiting the pathway further downstream. Here, we examined the effect of destabilizing HIF on ciliary mechanical properties by adding 30 *µ*M of the small molecule HIF inhibitor CAY10585 (19) to the cell culture media for 24 hours. Figure 3 presents our data.

### Experimental Error analysis

To demonstrate that our experimental results are valid, we provide some error analysis. The errors in our best-fit values result from a combination of random and systematic errors; systematic error will be discussed first.

#### Measurement of trap stiffness

We checked for any systematic error or variation in the measured trap stiffness as a function of trap height. This check was performed in case the QPD signal varied with optical path length. Using the microscope z-axis controller, we trapped microspheres at different heights above the glass slide and computed the asymptotic value of the MSD. Our data is shown in Figure 4. Our data shows that the trap stiffness varies very weakly with trap height and can be neglected. Thus, we have confidence that the length dependence of ciliary flexural rigidity is not an artifact.

**Figure 4:**
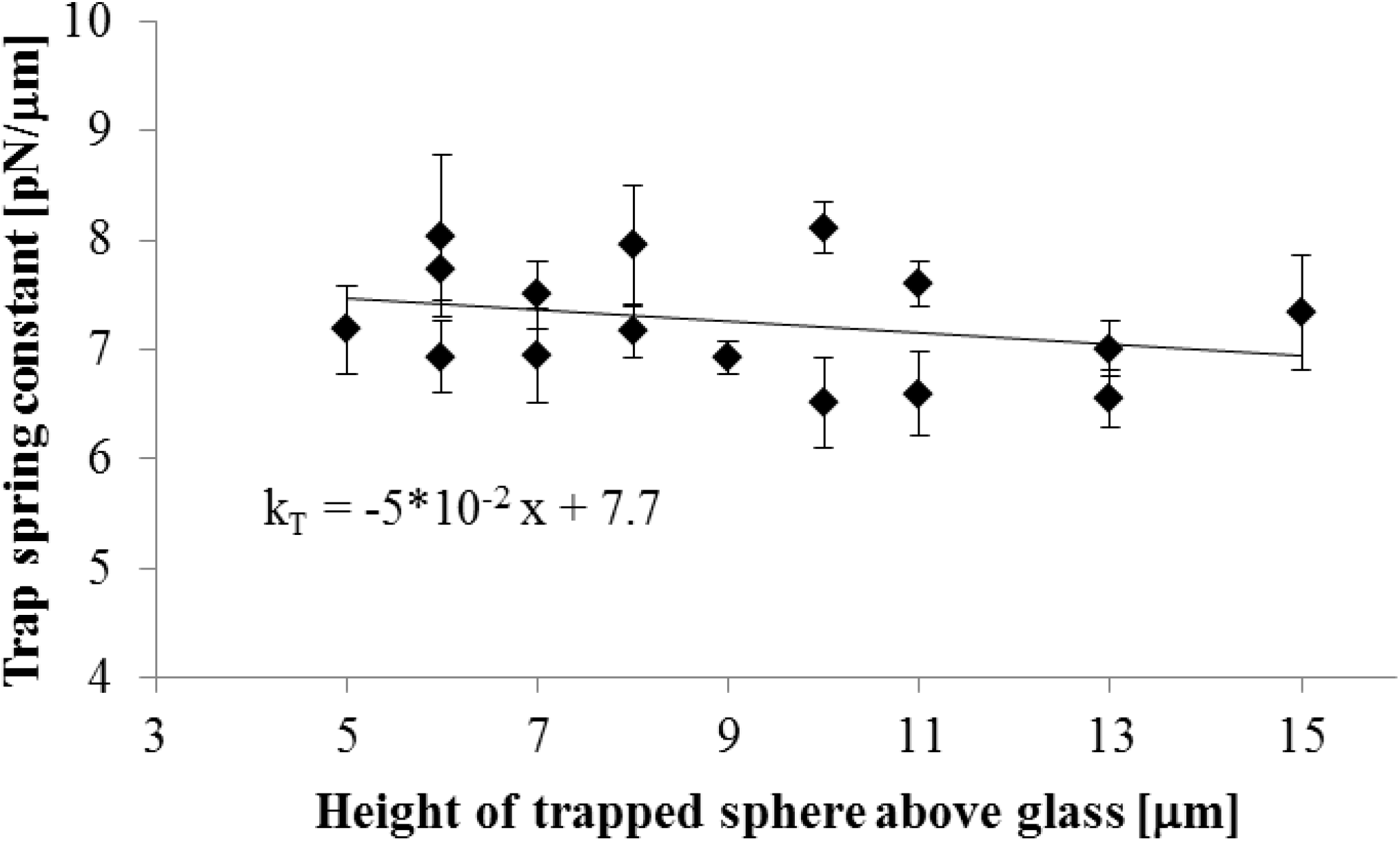
asymptotic MSD of optically trapped microspheres

#### Sources of random error

The primary sources of independent random error are: calculation of the MSD asymptote, measurement of cilium length, and experiment-to-experiment variability of the trap stiffness. As discussed in (22), the variability of MSD asymptote is reduced due to processing independent data subsets, the variation of this parameter is typically 10%. The variation due to length measurement varies with length because our uncertainty is fixed by the depth of focus of the objective lens, *δL* = ±0.3 *µ*m. Uncertainties due to experimental variations in the trap stiffness are caused by random variations of many parameters: optical alignment (including laser pointing stability, etc.), relative index of refraction between trapped object and surrounding fluid medium, diameter of primary cilium, potential mechanical disturbances to the optical bench, and so on. Because these collective uncertainties would simply result in random uncertainty in the calculated MSD asymptote, we may estimate the total random error of the MSD to be on the order of 10%.

### Theoretical Analysis

In this section, we provide the results of several approaches that attempt to explain the apparent length dependence of EI_*eff*_.

#### Structural model of a primary cilium

Following (14, 15, 17), we initially modeled the cilium as a uniform cantilevered beam of length ‘L’ subject to external loading. This ‘elastica’ model for a primary cilium treats the structure as a homogeneous isotropic flexible cylindrical beam with a hemispherical endcap, constrained at the basal end and free to move at the distal end. This model has been used for a wide variety of systems, including filamentous biopolymers (34), atomic force microscope tips (35), and cellular protrusions including flagella (36), stereocilia (37), glycocalyx (38) and actin brush border (39).

Because primary cilia, unlike flagella or motile cilia, do not actively generate internal forces, we may model the primary cilium as a passive beam: there are no forces and/or moments generated within the beam. Because the slenderness (length/diameter) of the cilium is large, we may also neglect both rotatory inertia and transverse shear and approximately describe the cilium shape in terms of a 1-D object, the so-called neutral axis (30).

In the Supplementary section, We present two variations of this model: the homogeneous ‘classical’ cantilever and the homogeneous ‘generalized’ cantilever’. Compared to the classical model, the generalized cantilever has a different boundary condition at the fixed end.

#### Model verification

The two models can be experimentally verified or discarded by use of an optical trap. We will demonstrate that our measurements can determine an effective cilium spring constant *k*_*cilium*_, which for each model is given by:

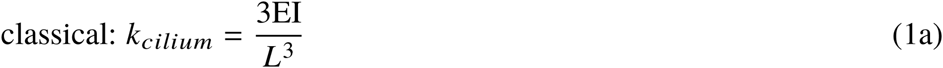

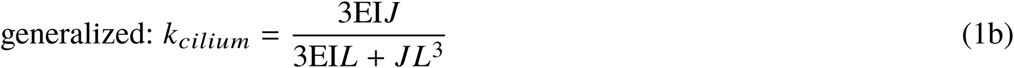

In both cases, the quantity *L*^3^*k*_*cilium*_ is equivalent to an effective bending modulus EI_*eff*_ for a cilium of length ‘L’. That is to say, if a cilium can be modeled as a homogeneous, isotropic material then our measurements should result in EI_*eff*_ ∝ *L*^3^. However, our data (see Figure 2) provides evidence that the effective bending modulus of a cilium EI_*eff*_ ∝ *L*^4^.

Because neither set of boundary conditions result in predictions that match measured data, we seek to identify other mechanisms. One potential solution may be to allow the bending modulus to vary with length EI = EI(s), however the Euler-Bernoulli equation itself is then modified, meaning our use of equations 1a or 1b to fit EI_*eff*_ are not appropriate. Unfortunately, the general equation that results from allowing EI = EI(s) is very cumbersome, not obviously solvable, and thus presented as Supplementary material. Importantly, because the ciliary axoneme has a constant cross section and essentially a uniform composition along the length, it is unclear what could cause a spatially-varying bending modulus.

Consequently, we will now carefully examine the stochastic model for optically trapped objects immersed in a viscous medium as it applies to a cilium and focus specifically on viscous drag as a potential way to reconcile our experimental results with prior models.

### Stochastic model of an optically trapped cilium

Application of an optical trap to a cantilevered beam introduces at least two complications as compared to a trapped free particle, which we model in the Supplementary section. First, hydrodynamic (viscous) forces act along the entire length of the cantilever and not just the trapped end. Second, although the distal end of the cantilever can ‘freely’ respond to the applied trap force, the constrained end introduces a restoring force, embodied by the cantilever spring constant (equations 1a, 1b). In equilibrium, the tip of the cilium will be not be located in the center of the trap but displaced from the center by a distance ‘x’ given by the balance of trapping and elastic restoring forces. We denote the distance from the unbent cilium axis to the trap axis as ‘d’.

#### Simplified model of the trapped cilium

We first consider a simplified model of the cilium- the axoneme is subject to viscous forces but the viscous damping coefficient *γ* remains unspecified. Figure 1 represents this simplified model of the primary cilium.

From Figure 1, we have a modified Langevin-type equation:

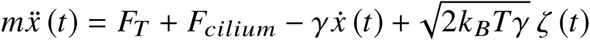

Where 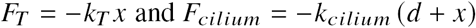.

Rearranging, we obtain our equation of motion:

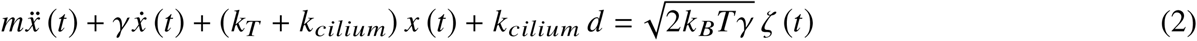

A key point to emphasize here is that while k_*T*_ should be the same for every trapped cilium, *k*_*cilium*_ may vary from cilium to cilium. The Fourier transform of Equation 2 is:

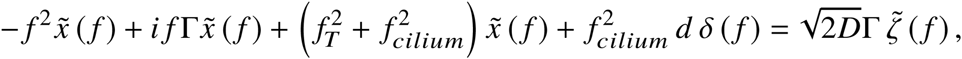

where *δ* () is Dirac’s delta function. Defining 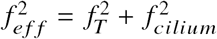 we obtain

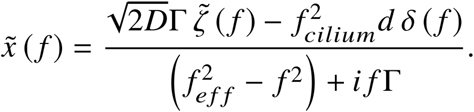

Next to find the MSD of the cilium tip, we calculate the PSD first, noting that the term 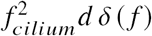 is not stochastic but rather a constant offset and can be dropped:

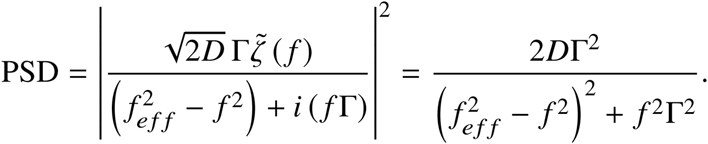

We can obtain the MSD of the trapped cilium tip as before:

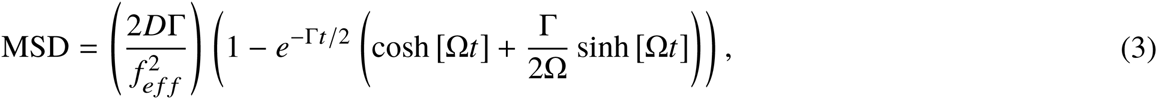

where 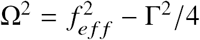 and as shown in the Supplementary section, the asymptotic limit as 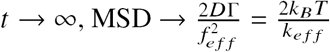.

The results obtained now provide useful information regarding the overall parameterization of the MSD in terms of the mechanical properties of the trapped cilium. This can be shown by combining the asymptotic limit of the MSD with equations 1a or 1b:

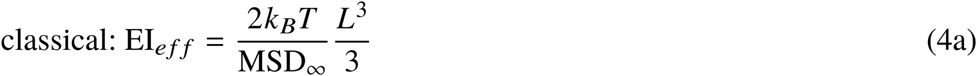

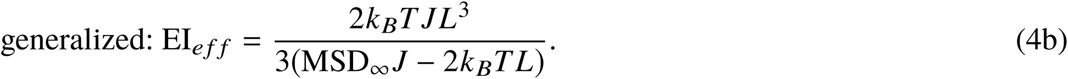

Again, we find that in either case, EI_*eff*_ ∝ *L*^3^, in conflict with our measurements. In the Supplementary material, we show that even when the drag coefficient *γ* is correctly calculated for a cylinder of length ‘L’, there is no change to the asymptotic form of the MSD. Our computational coarse-grained model further confirms that the hydrodynamic viscous force along the entire cilium can more simply be represented as an ‘effective’ force localized to the trapped end. Consequently, accounting for viscous drag does not ”rescue” either homogeneous cantilever model.

Taken together, we conclude that our experimental results are not due to an experimental artifact, systematic error, or random error. Furthermore, neither incorporation of different boundary conditions, nor accounting for viscous drag, nor spatial variations in EI result in accurate modeling of our results. We must now consider other ways to alter the structural model of primary cilia in ways that still preserve prior results, so we turn to computational and analytical methods to more efficiently explore the parameter space.

### Improved analytical shell model of cilium structure

Our next approach is to refine the structural model of primary cilia. Rather than considering the ciliary axoneme as effectively homogeneous, following (18) and (40) we now treat each axonemal microtubule as an orthotropic elastic shell with wall thickness h = 2.7 nm and middle radius R = 12.5 nm.

The essential result of this model (see Supplementary materials) is that the persistence length of the microtubules, which is proportional to EI, depends on the aspect ratio *α* = *L*/*R*:

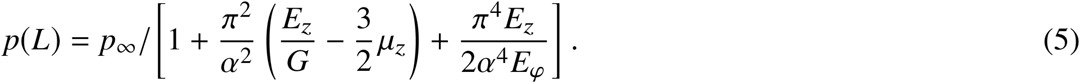

Using our prior results from Eqn 1a and the worm-like chain (WLC) model (32) EI_*eff*_ = *p*(*L*)*k*_*B*_*T*,

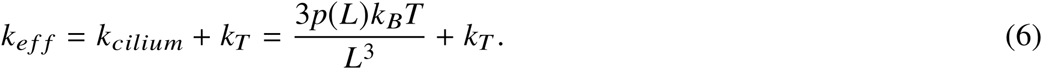

Modeling a cilium in terms of a bundle of elastic shells results in a significant departure from homogeneous models; while homogeneous models predict EI_*eff*_ ∝ *L*^3^, the shell model predicts EI_*eff*_ ∝ *L*^4^, matching the data trend.

### Basal Body fluctuations

We now consider the role of basal body dynamics on ciliary motion (21). By applying our coarse-grained DPD model, we calculated the mean-squared displacement (MSD) of the tip held within an optical trap when the entire cell is within a hydrodynamic thermal bath consisting of DPD particles of water held at 39°C. We found that the MSD of the tip always increases with cilium length when we use either a constant persistence length for the microtubules or use the length dependent persistence length from Eq. 30. Since we model the basal body as a rigid body described by the Langevin equation and model the remaining cellular components within the DPD framework, our model allows us to assign different temperatures for the Langevin equation (basal body) and the hydrodynamic bath. This approach, using a higher temperature for the basal body, models actin-driven ”active fluctuations” of the basal body. Our model predicts the MSD of the tip decreases with the cilium length for either constant or length-dependent microtubule persistence lengths, which is qualitatively consistent with our experimental observations. Informed by these simulation results, we now derive a simplified analytical model for the induced fluctuations of the cilium tip caused by active fluctuations of the basal body.

The most simple way to incorporate stochastic motion of the basal body is to alter eqn 2:

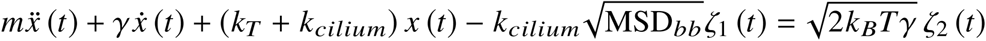

Where we have allowed the basal body to undergo active fluctuations with a mean-squared displacement MSD_*bb*_ and we use two different (uncorrelated) white-noise processes *ζ*_1_ and *ζ*_2_. Going through the usual calculations, we obtain *MSD*_*∞*_:

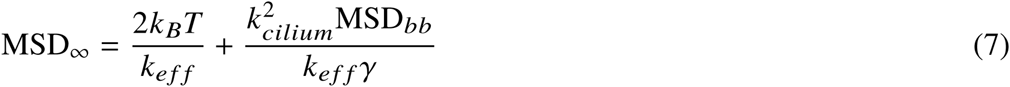

Importantly, because *k*_*cilium*_ appears in both terms but *MSD*_*bb*_ in only one, there is a possiblity to independently alter *k*_*cilium*_ and *MSD*_*bb*_, providing multiple biochemical pathways to probe the ciliary flow response.

### Parameter Estimation

It is important to realize that our analytical model only has a few free parameters; material properties of untreated microtubules have been independently measured, for example the ratio 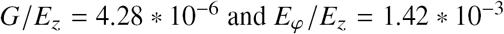(18). Regarding the two different homogeneous models, our computational model showed that axonemal bending dominates over the rotations of the basal body, we may set the rotational stiffness of the basal body *J* = ∞, making the models equivalent.

We performed a least-squares fit of *k*_*eff*_ based on our measured MSD_∞_ to obtain estimates for the only two free parameters contained in eqns 5, 6 and 7: *k*_*T*_ and *MSD*_*bb*_ / *γ*_0_ (*γ*_0_ = *γ*/*L*). Following (21), we considered the axonemal microtubules as mechanically uncoupled. The result is shown is Figure 5 corresponds to values of MSD_*bb*_ / *γ*_0_ = 5.89 * 10^−6^ *μ*m^4^ /*p*_*N*_, and *k*_*T*_ = 15.5 pN *μm*. In the Supplementary section, we provide an estimate for *γ*_0_ = 0.5*pN/μm*, so that our model predicts basal body fluctuations on the order of 2 nm.

**Figure 5:**
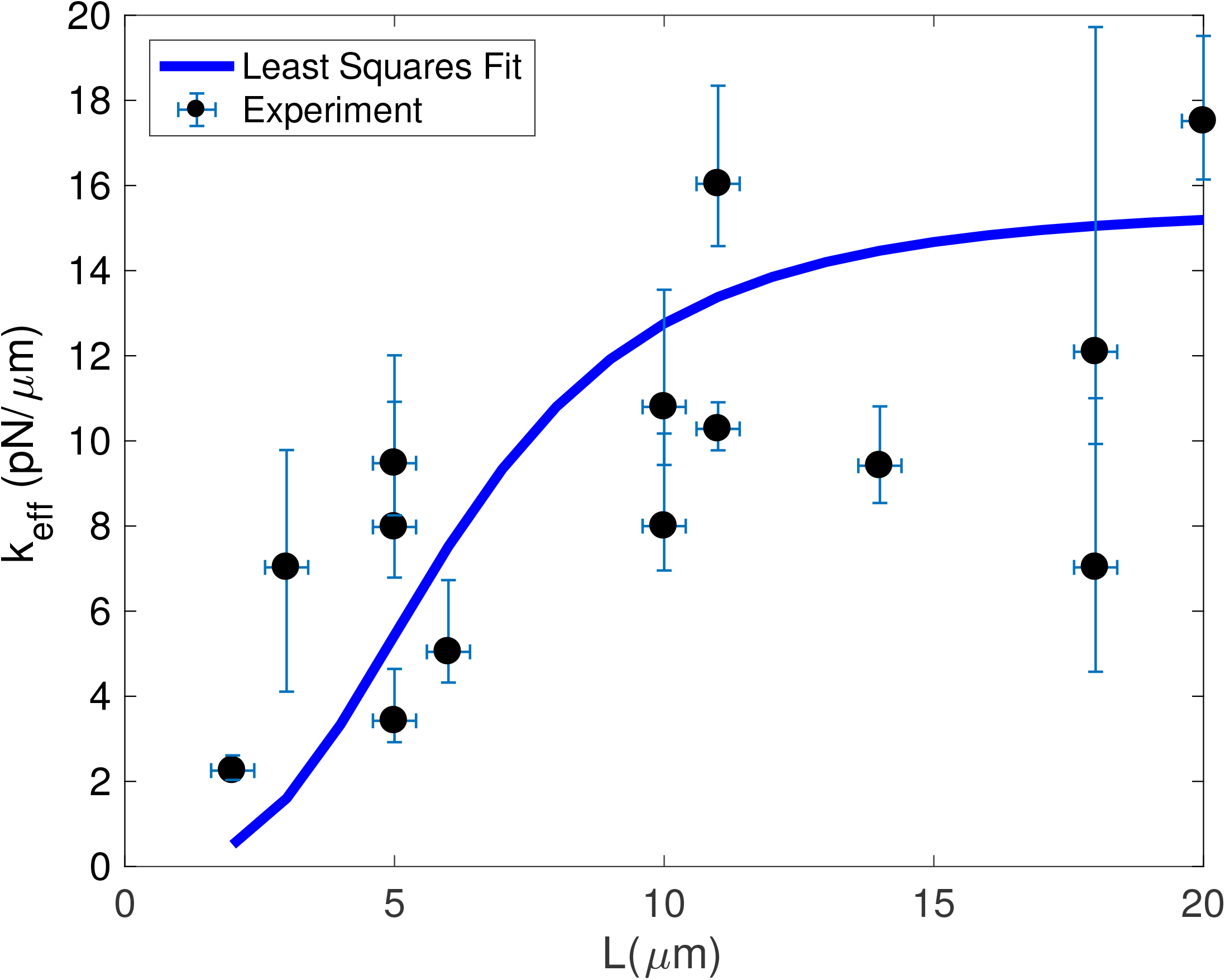
Best-fit results for *k*_*eff*_ as a function of cilium length, untreated cilia

Performing the same least-squares fit on our datasets of treated cilia, we obtain *G*/*E*_*z*_ = 4.04 * 10^−7^ and MSD_*bb*_/*γ*_0_ = 1.88 * 10^−5^/*μm*^4^/*pN* for cells treated with Taxol, while for HIF-inhibitor treated cells we obtain *G/E*_*z*_ = 1.39 * 10^−6^ and MSD_*bb*_/*γ*_0_ = 3.192 * 10^−7^ *μm*^4^/*pN.* This provides crucial information about the effect of the two drugs; Taxol directly binds to tubulin and primarily impacts the mechanical properties of the axoneme (32) while suppressing HIF pathways primarily decreases actin fluctuations of the basal body (21). Consequently, our data also demonstrate the ability to target different components of the overall mechanical response of a cilium.

## CONCLUSION

We have attempted to demonstrate that a systematic treatment of end-loaded cantilevers immersed in a viscous medium can be applied to a biological system of high relevance: the primary cilium. Careful treatment demonstrates that (1) in contrast to previous efforts, cilia cannot be modeled as a homogeneous cantilever but rather as composed of orthotropic shells, and (2) fluctuations of the basal body are an essential component of ciliary mechanics. In addition, we have demonstrated that our analytical method is well-matched to a class of experimental techniques, including optical trapping but also other related approaches (magnetic trapping, for example) that apply localized forces to the cilium. We explored how the mechanical response of the microcantilever relates to the underlying mechanical properties, and identified the basal end of the primary cilium as a site of high interest, both because the structural properties are largely unknown but also because the basal end could be the site of initiation of the mechanosensation response.

We have found, for the first time, evidence supporting a length-dependent effective bending modulus. It is possible that our data can help explain the persistent uncertainty in measured values of the ciliary bending modulus; in particular the finding (17) that longer cilia apparently have a higher apparent bending modulus. Finally, we presented data demonstrating the impact of various pharmacologic treatments on the effective bending modulus of cilia.

Our results begin to address a possible relationship between (regulated) cilium length and mechanical response to fluid flow. Regarding our hypothesis that flexural rigidity may be involved in the regulation of cilium length, the existing literature is inconclusive. One issue is that in the absence of fluid flow stimulation, cilium length is often heterogeneous (41), another is that fixation techniques damage the cilium (42). There is a tentative connection between increased cilium length and HIF stabilization (33) and we have recently published data showing that HIF stabilization results in more flexible cilia (20), so our hypothesis is reasonable. The work presented here should provide an increased ability to study our hypothesis.

It should be mentioned that in contrast to other reports, our results were obtained without application of fluid flow to deform the cilium. This is significant given Stokes’ paradox- small experimental uncertainties in the local flow velocity will have large effects on the applied shear stress and resultant deformation, potentially creating systematic parameter estimation error. Our approach avoids the need to characterize the local flow field and so is not impacted by that source of error.

Another avenue for extension of our results lie with the boundary conditions at the basal end, that is, how to account for the basal body. A growing body of results (12, 13, 15, 21) focus on the basal body both as a modifier of the mechanical response and as some sort of ‘gate’ regulating the transport of materials in and out of the cilioplasm, thus the basal body potentially serves as the site of cellular flow sensing. We have demonstrated the possibility of investigating basal body dynamics by observing ciliary dynamics.

## AUTHOR CONTRIBUTIONS

AR designed the research. JF and AR performed the experiments. ZF and ZP applied the shell model; AR, ZP, and Y-N Y analyzed the data. All authors contributed equally to writing the article.

## ACKNOWLEDGMENTS

AR’s research was supported by Dr. John Vitullo’s Pilot and Bridge Funding Program at the Center for Gene Regulation in Health and Disease (GRHD) (awarded to AR). YNY’s work was supported by a faculty seed grant from NJIT.

## SUPPLEMENTARY MATERIAL

### Continuum models of the primary cilium

#### Classical cantilevered beam

The time-dependent shape Y(s,t) of the neutral axis of a cilium is given by the linearized Euler-Bernoulli law for pure bending (see, for example, (43)):

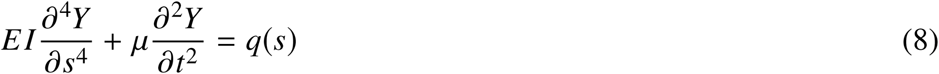

Where *q*(*s*) is an externally applied distributed load (force per length), ‘*μ*’ is the mass per unit length, ‘E’ is the Young’s modulus, ‘I’ the area moment of inertia (for a cylinder of radius a, I =*π*a^4^/4) and EI together referred to as the flexural rigidity or bending modulus, having units of Force*area. Measurements of the flexural rigidity of primary cilia vary, but generally cluster between EI = 10-50 pN·*μ*m^2^ (14, 16, 17, 21). We consider the cilium to be inextensible, and all deformations are considered transverse to the neutral axis. For simplicity, we reduce the full three-dimensional range of motion to deformations within a plane.

To solve the Euler-Bernoulli equation, we require boundary conditions for both the free and constrained ends. At the free (distal) end, the cilium responds to an applied load:

1. the bending moment vanishes:

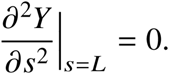

2. The free end is subject to shear from an applied end load, for example from the optical trap ‘*F*_*T*_’:

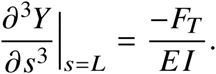

and the constrained end has a ‘built in’ support:

3. The constrained end cannot move: Y(0)=0.

4. The constrained end is ‘built in’:

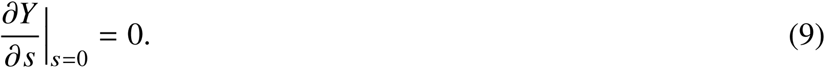

The classical cantilever has static solutions:

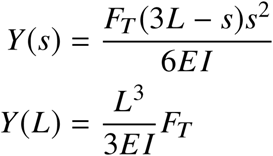

#### Generalized cantilevered beam

Typically, the constrained end of cantilevered beams is classified as ‘free’, ‘clamped’ or ‘hinged’, depending on how the cantilever is attached to a substrate. The most general case of an end connected to a translational and rotatory spring, linear and torsional damper, and mass with rotational inertia is covered in (43). While boundary conditions for the basal attachment of cilia are not yet fully known, preliminary results (15, 16, 21) have shown that the fixed end is neither free, clamped nor hinged. One current model treats the fixed end as constrained by a nonlinear rotatory spring obeying a force law F(*θ*) = J*θ* + *αθ*^2^ (15, 16), where ‘J’ is the linear (Hookean) spring constant and ‘*α*’ the nonlinear coefficient, while another incorporates a viscoelastic rotatory spring (21). The more general boundary condition replacing the fourth boundary condition (equation 9) is:

4. The constrained end is attached to a mass with rotational inertia *I*_*m*_ by a nonlinear torsional spring and linear torsional damper (coefficient ‘c’):

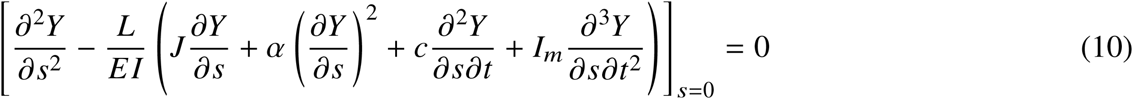

With this boundary condition, the static solution to the Euler-Bernoulli equation is found to be:

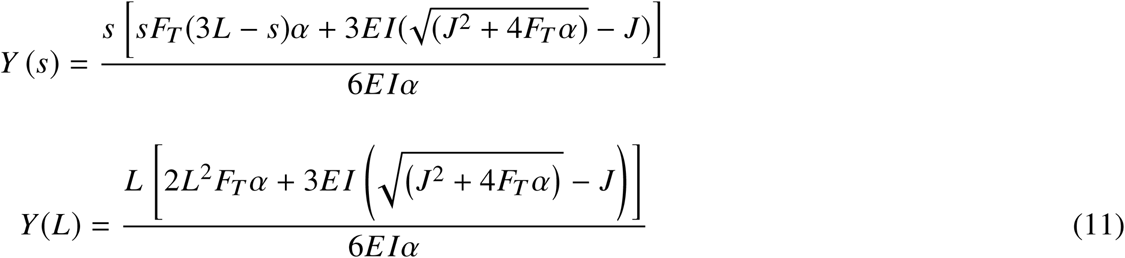

The solution is constrained by requiring L > 0, J > 0, and EI > 0.

#### Beam bending equation with spatially varying bending modulus

For the linear theory of pure bending, points along the neutral line defined by the axial coordinate ‘s’ will be strained by an amount

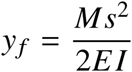

where M is the bending moment and I the ‘second moment of area’. If we allow *EI* = *EI*(s), meaning the quantity EI may collectively vary with s, then along the neutral axis:

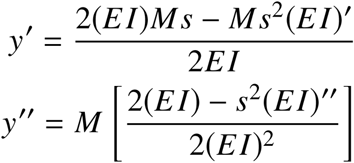

where 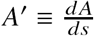. For static deflections generated by a distributed load *w*(*s*)

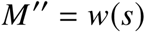

which, after substitution results in:

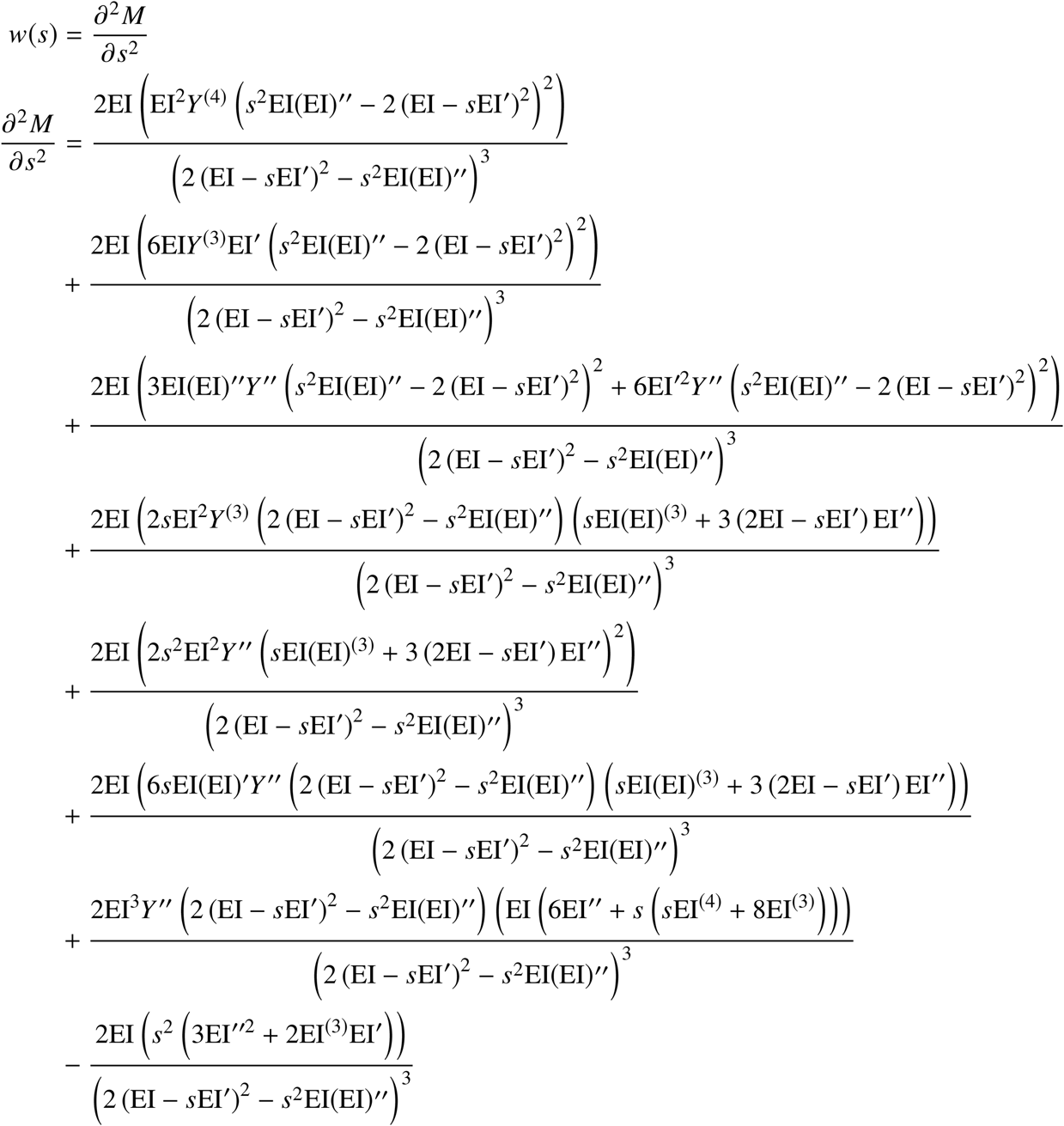

This expression reduces to the usual Euler-Bernoulli equation if EI is constant.

### Stochastic model of an optically trapped object

The Langevin approach provides a convenient formalism to analyze the stochastic motion of a particle that undergoes Brownian motion while held within an optical trap, so-called constrained Brownian motion. Following (44), we analyze the motion of a particle of mass ‘m’ within a viscous fluid (viscous damping *γ*) experiencing Brownian stochastic forces *F*_*B*_ while suspended within an optical trap with trapping force −*k*_*T*_ *x*.

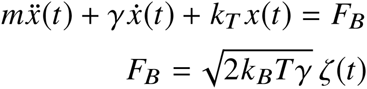

*ζ*(*t*) is a normalized white-noise process. Performing the Fourier transform on this equation, using the definitions: trap corner frequency 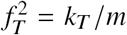; normalized viscous drag 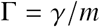, diffusion coefficient 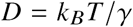, and the standard notation for Fourier transform pairs:

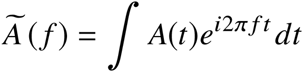

Gives

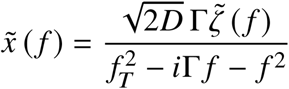

The power spectral density (PSD) is given by the square modulus: PSD = 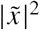.

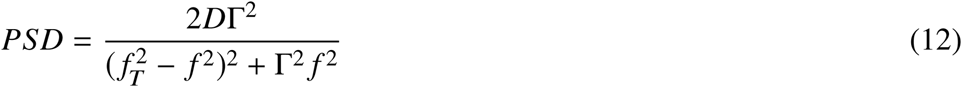

As mentioned previously, precision calibration of optical traps typically analyze the PSD (26, 45), but that precision requires knowledge of the viscous damping Γ. This is important because in contrast to a sphere, the viscous drag acting on a cylindrical object is non-trivial. Our analysis uses the MSD of a trapped object rather than the PSD because analytic expressions for the viscous damping are not required. By the Wiener-Khinchin theorem, the PSD is equal to the Fourier transform of the MSD. Because the MSD is a real-valued function, the cosine-Fourier transform is used. In our case, trapped cilia immersed in a viscous fluid can be described as an overdamped system (*f*_*T*_ < Γ/2):

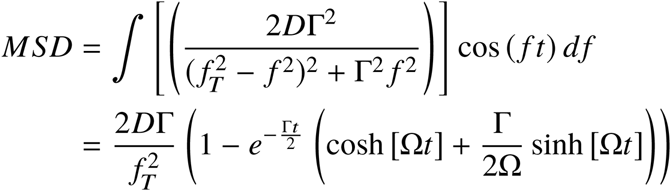

where 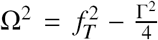. Experimentally, it is useful to consider the limit of the MSD as *t* → ∞. Expanding the hyperbolic functions and linearizing Ω. by making use of Γ ≫ *2f*_*T*_, we obtain:

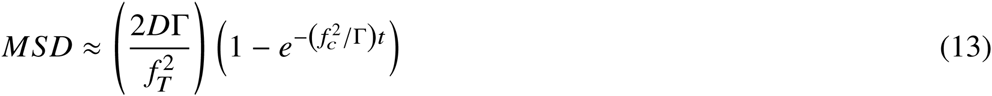

Thus, as *t* → ∞, 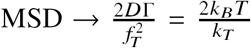. That is, the trap spring constant can be calculated by 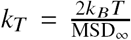, where we denote the long-time asymptotic value of the MSD by MSD_∞_. As we show, this form of MSD_∞_ is very robust to more detailed modeling.

### Exact results for a optically trapped homogeneous cantilever immersed in a viscous medium

Incorporation of hydrodynamic forces along the ciliary axoneme is complicated because there is no solution to the Navier-Stokes system of equations for flow around a circle (Stokes Paradox) (46). The approach (35) used here considers the fluid drag to act as an (additional) effective mass, and we must resolve the Stokes paradox.

Beginning with the Fourier transformed scaled time-dependent Euler-Bernoulli equation (eqn 8), we account for hydrodynamic viscous drag *F*_*Hydro*_ by a fluid with density *ρ*, viscosity *η*, and an applied driving force *F*_*d*_:

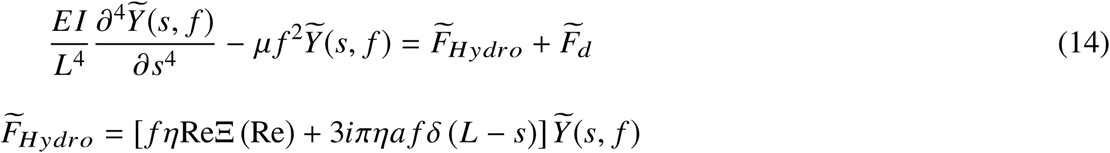

Re is the Reynolds number 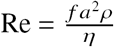. For primary cilia, Re ≪ 1. The hydrodynamic drag terms correspond to flow around a cylindrical axoneme and flow around the hemispherical tip, respectively.

In a sense, the drag coefficient Ξ (Re) summarizes the Stokes paradox. Detailed derivations resolving the Stokes paradox can be found elsewhere (46, 47) so we only summarize the results, originally derived by Oseen and Lamb (48). When the cylinder axis is perpedicular to the fluid velocity, the first terms of the hydrodynamic drag coefficient are given by (49):

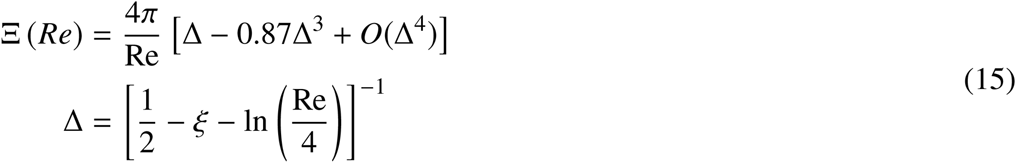

where *ξ* is Euler’s constant (*ξ* = 0.577…). The hydrodynamic drag force *F*_*Hydro*_ → 0 as Re → 0, but importantly, the drag coefficient diverges: Ξ (Re) → Re^−1^ as Re → 0. This divergence has the effect of drastically decreasing the resonant frequency of a cilium, from approximately 100 kHz in vacuum to 50 Hz in an aqueous medium (15) and greatly complicates use of the PSD to analyze trapped cilia.

The driving force *F*_*d*_ consists of two terms: Brownian forces F_*B*_ and the optical trap F_*T*_. We assume the optical trap applies a force only to the tip of the cilium *F*_*T*_ = *F*_0_*δ*(*s* – *L*). For our trap the beam waist is 0.3 *μ*m and Reyleigh length is 0.4 *μ*m, justifying the use of a localized applied force.

#### Calculation of the PSD

Constructing the eigenfunction mode expansion (index ‘n’) of the cantilever (50)

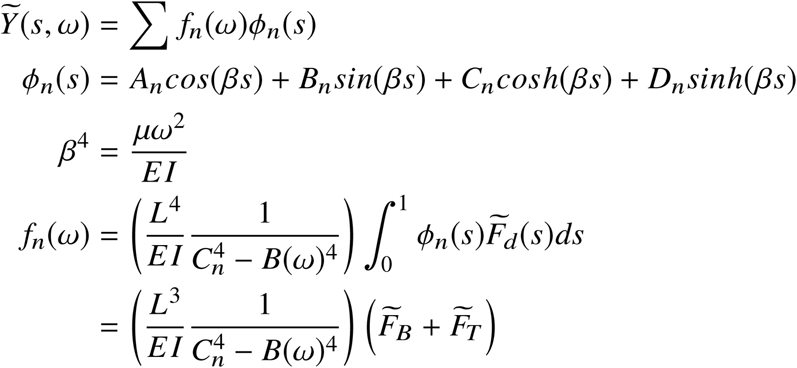

Where 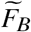 is the Brownian force integrated along the cilium length and 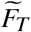 the trap force, which is localized to the cilium tip:

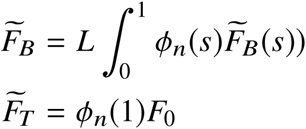

We also defined

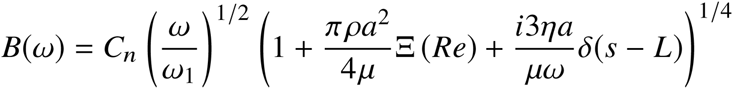

*C*_*n*_ = 1.875, 4.694, 7.854, 10.995… are roots of the eigenvalue characteristic equation and 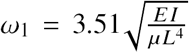 is the lowest eigenvalue for oscillation in an inviscid medium. For additional details, please see Ref. (35).

The eigenmode expansion simplifies considerably for trapped cilia when evaluated at the distal tip. In this case, a single mode (n = 1) is dominant and the Brownian force is uncorrelated with the (static) trap force:

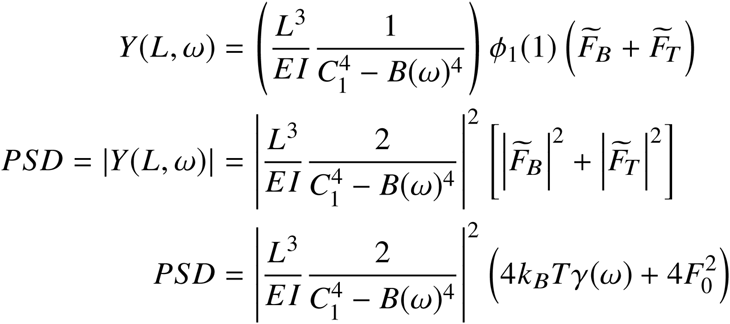

Where *γ*(*ω*) is the total damping on the entire cilium. After a considerable amount of algebra, we obtain:

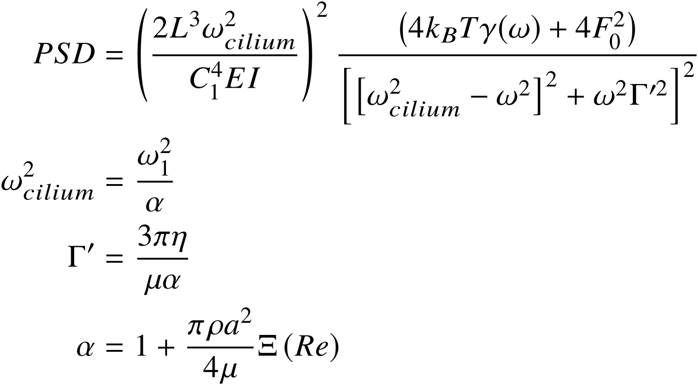

Using prior definitions 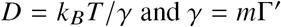 and noting that the trap force is a constant offset, this simplifies to

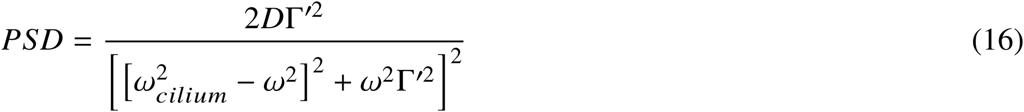

with the damping 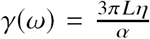, when *Re* << 1 this simplifies to 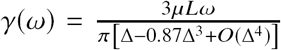. Again, as *t → ∞,* MSD 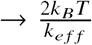 and we obtain the cilium spring constant based on MSD_∞_. Note, the damping coefficient still does not appear in MSD_∞_. Using representative values for the fluid viscosity, mass density of a cilium, and Reynolds number (15), we estimate the parameter 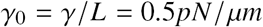.

### Setup of the computational coarse-grained model of the primary cilium

As shown in figure 6, we explicitly model the 9 vertical microtubule doublets of the axoneme (covered by cilia membrane) and cytoplasmic microtubules (green) connecting basal body to the actin cortex (blue) and the lipid-bilayer (red). The basal body including the centrioles and surrounding pericentriolar material (PCM) was modeled as a rigid sphere, which can rotate under the constraints from connecting microtubules. The microtubules in the axoneme and the cytoplasmic microtubules are connected to the basal body as a clamped boundary condition. More importantly, the actin cortex and its interaction with microtubules were also explicitly modeled. The lipid-bilayer (red) and the actin cortex (blue) are modeled as two distinct components that couple together, with a two-dimensional triangulated network for the cortex. Details of this model can be found in (27).

**Figure 6:**
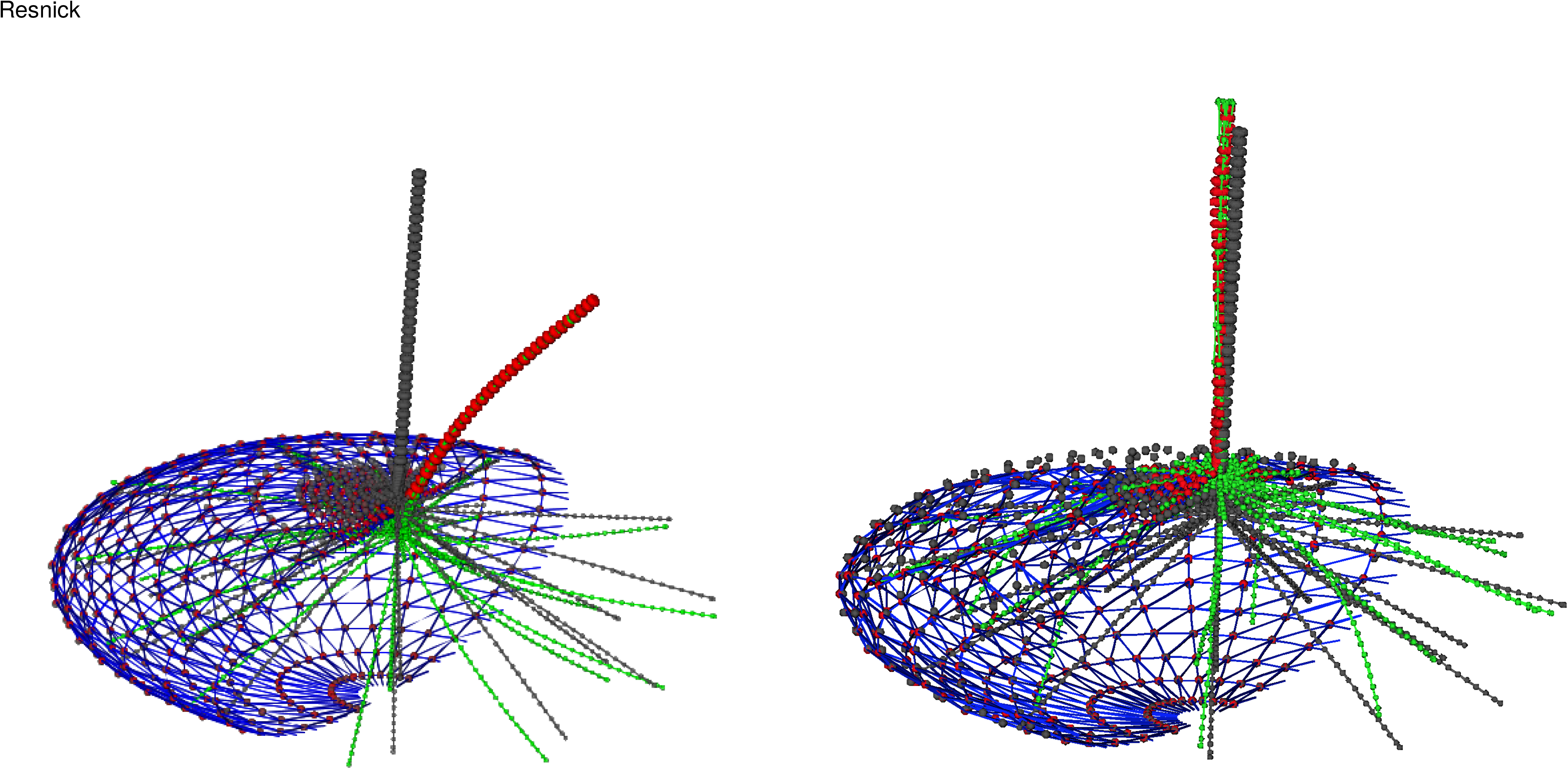
Coarse-grained simulations (only half of the cell membrane shown). Cilium shown in both untrapped/trapped configurations (left) and thermal fluctuations (right).

### Governing equation of an orthotropic cylindrical shell

In the cylindrical coordinate system (r,*φ*,z), the strains of the shell are related to the displacement fields as (refer to (18))

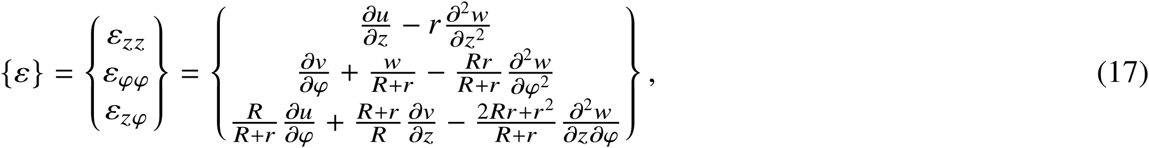

where *u*(*z, φ*), *v*(*z*, *φ*), *w*(*z, φ*) are displacements in z, *φ, r* directions. For orthotropic linear elastic materials, the stresses are related to the strains as

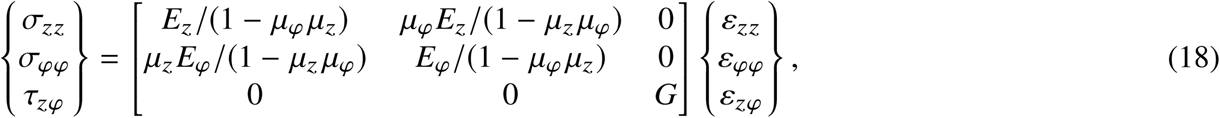

where *μ*_*α*_ are the Poisson ratios, *E*_*α*_ the Young’s moduli 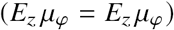, and G the shear modulus. The stress resultants are related to the stresses and displacement fields; referring to (18) we obtain expressions for the internal normal and shear forces and moments of a shell element undergoing small displacements:

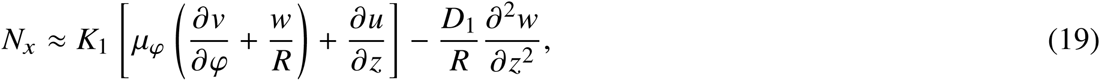

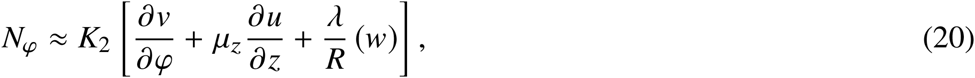

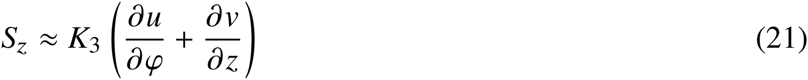

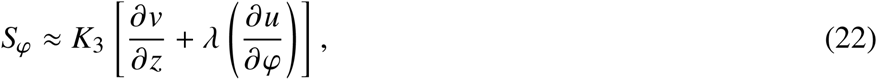

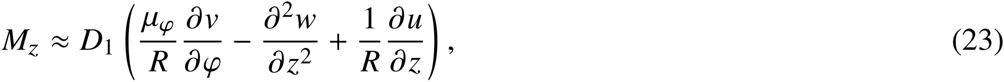

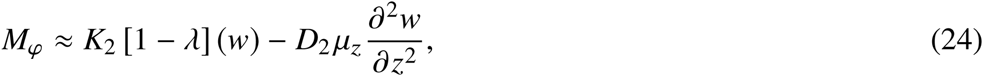

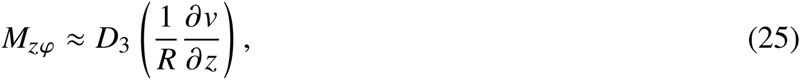

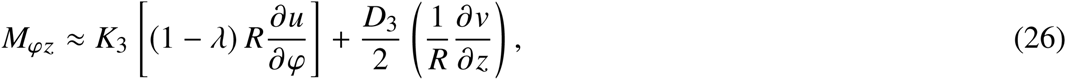

where the stretching stiffness coefficients 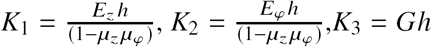; bending stiffness coefficients 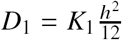, 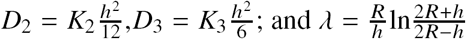; *h* is the shell (effective) thickness.

For the case of pure bending by a point end load ‘q’, the forces and moments obey the following set of differential equations:

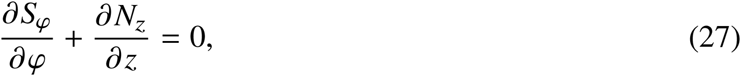

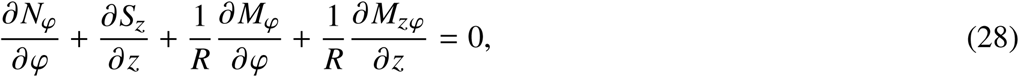

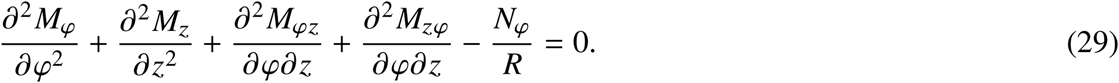

Inserting expressions for the stress resultants (Eqns. (19)-(26)) into these three equilibrium equations (Eqns. (27)-(29)), provides, after a lengthy calculation ((40)) the contour-length dependent persistence length when a single bending mode dominates (n=1):

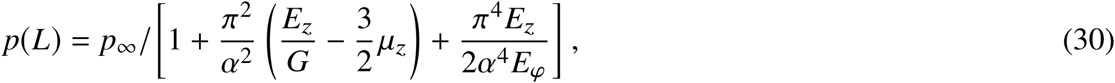

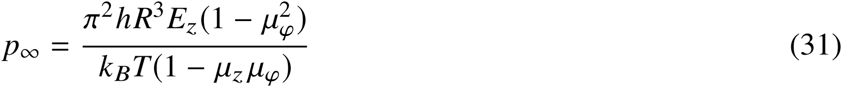

where *α = L*/*R* ≫ 1.

An online supplement to this article can be found by visiting BJ Online at http://www.biophysj.org.

## REFERENCES

1. Delling, M., P. G. Decaen, J. F. Doerner, S. Febvay, and D. E. Clapham, 2013. Primary cilia are specialized calcium signalling organelles. Nature 504:311–314.

2. Jin, X., A. M. Mohieldin, B. S. Muntean, J. A. Green, J. V. Shah, K. Mykytyn, and S. M. Nauli, 2013. Cilioplasm is a cellular compartment for calcium signaling in response to mechanical and chemical stimuli. Cell. Mol. Life Sci. Cellular and Molecular Life Sciences 71:2165–2178.

3. Resnick, A., 2010. Use of optical tweezers to probe epithelial mechanosensation. J. Biomed. Opt Journal of Biomedical Optics 15:015005.

4. Praetorius, H., and K. Spring, 2001. Bending the MDCK Cell Primary Cilium Increases Intracellular Calcium. Journal of Membrane Biology 184:71–79.

5. Lin, F., T. Hiesberger, K. Cordes, A. M. Sinclair, L. S. B. Goldstein, S. Somlo, and P. Igarashi, 2003. Kidney-specific inactivation of the KIF3A subunit of kinesin-II inhibits renal ciliogenesis and produces polycystic kidney disease. Proceedings of the National Academy of Sciences 100:5286–5291.

6. Ma, M., X. Tian, P. Igarashi, G. J. Pazour, and S. Somlo, 2013. Loss of cilia suppresses cyst growth in genetic models of autosomal dominant polycystic kidney disease. Nature Genetics Nat Genet 45:1004–1012.

7. Delling, M., A. A. Indzhykulian, X. Liu, Y. Li, T. Xie, D. P. Corey, and D. E. Clapham, 2016. Primary cilia are not calcium-responsive mechanosensors. Nature 531:656–60.

8. Bloodgood, R. A., 1990. Ciliary and flagellar membranes. Plenum Press, New York.

9. Nauli, S. M., F. J. Alenghat, Y. Luo, E. Williams, P. Vassilev, X. Li, A. E. H. Elia, W. Lu, E. M. Brown, S. J. Quinn, and et al., 2003. Polycystins 1 and 2 mediate mechanosensation in the primary cilium of kidney cells. Nature Genetics Nat Genet 33:129–137.

10. Garcia, r., G., and J. F. Reiter, 2016. A primer on the mouse basal body. Cilia 5:17.

11. Mofrad, M. R. K., and R. D. Kamm, 2010. Cellular mechanotransduction: diverse perspectives from molecules to tissues. Cambridge University Press, Cambridge; New York.

12. Hu, Q., and W. J. Nelson, 2011. Ciliary diffusion barrier: the gatekeeper for the primary cilium compartment. Cytoskeleton (Hoboken) 68:313–24.

13. Lin, Y. C., P. Niewiadomski, B. Lin, H. Nakamura, S. C. Phua, J. Jiao, A. Levchenko, T. Inoue, and R. Rohatgi, 2013. Chemically inducible diffusion trap at cilia reveals molecular sieve-like barrier. Nat Chem Biol 9:437–43.

14. Schwartz, E. A., M. L. Leonard, R. Bizios, and S. S. Bowser, 1997. Analysis and modeling of the primary cilium bending response to fluid shear. American Journal of Physiology - Renal Physiology 272:F132–F138.

15. Resnick, A., 2015. Mechanical Properties of a Primary Cilium As Measured by Resonant Oscillation. Biophysical Journal 109:18–25.

16. Young, Y.-N., M. Downs, and C. Jacobs, 2012. Dynamics of the Primary Cilium in Shear Flow. Biophysical Journal 103:629–639.

17. Downs, M. E., A. M. Nguyen, F. A. Herzog, D. A. Hoey, and C. R. Jacobs, 2014. An experimental and computational analysis of primary cilia deflection under fluid flow. Comput Methods Biomech Biomed Engin 17:2–10.

18. Liu, X., Y. Zhou, H. Gao, and J. Wang, 2012. Anomalous Flexural Behaviors of Microtubules. Biophysical Journal 102:1793–1803. https://doi.org/10.1016%2Fj.bpj.2012.02.046.

19. Lee, K., J. H. Lee, S. K. Boovanahalli, Y. Jin, M. Lee, X. Jin, J. H. Kim, Y. S. Hong, and J. J. Lee, 2007. (Aryloxyacetylamino)benzoic acid analogues: A new class of hypoxia-inducible factor-1 inhibitors. J Med Chem 50:1675–84.

20. Resnick, A., 2016. HIF Stabilization Weakens Primary Cilia. PLoS One 11:e0165907.

21. Battle, C., C. M. Ott, D. T. Burnette, J. Lippincott-Schwartz, and C. F. Schmidt, 2015. Intracellular and extracellular forces drive primary cilia movement. Proc Natl Acad Sci U S A 112:1410–1415.

22. Glaser, J., D. Hoeprich, and A. Resnick, 2014. Near real-time measurement of forces applied by an optical trap to a rigid cylindrical object. Optical Engineering 53.

23. Leitz, G., E. Fallman, S. Tuck, and O. Axner, 2002. Stress response in Caenorhabditis elegans caused by optical tweezers: wavelength, power, and time dependence. Biophys J 82:2224–31.

24. Neuman, K. C., E. H. Chadd, G. F. Liou, K. Bergman, and S. M. Block, 1999. Characterization of photodamage to Escherichia coli in optical traps. Biophys J 77:2856–63.

25. Jones, P., O. Marag, and G. Volpe, 2015. Optical Tweezers: Principles and Applications. Cambridge University Press.

26. Norrelykke, S. F., and H. Flyvbjerg, 2011. Harmonic oscillator in heat bath: Exact simulation of time-lapse-recorded data and exact analytical benchmark statistics. Physical Review E 83.

27. Peng, Z., X. Li, I. V. Pivkin, M. Dao, E. Karniadakis, and S. Suresh, 2013. Lipid bilayer and cytoskeletal interactions in a red blood cell. Proc. Nat. Acad. Sci. 110:13356–13361.

28. Groot, R. D., and P. B. Warren, 1997. Dissipative particle dynamics: Bridging the gap between atomistic and mesoscopic simulation. J. Chem. Phys. 107:4423–4435.

29. Hoogerbrugge, P. J., and J. M. V. A. Koelman, 1992. Simulating microscopic hydrodynamic phenomena with dissipative particle dynamics. Europhys. Lett. 19:155–160.

30. Mickey, B., 1995. Rigidity of microtubules is increased by stabilizing agents. The Journal of Cell Biology 130:909–917.

31. Felgner, H., R. Frank, and M. Schliwa, 1996. Flexural rigidity of microtubules measured with the use of optical tweezers. Journal of Cell Science 109 (Pt 2):509–516.

32. Sept, D., and F. C. MacKintosh, 2010. Microtubule Elasticity: Connecting All-Atom Simulations with Continuum Mechanics. Physical Review Letters 104.

33. Verghese, E., J. Zhuang, D. Saiti, S. D. Ricardo, and J. A. Deane, 2011. In vitro investigation of renal epithelial injury suggests that primary cilium length is regulated by hypoxia-inducible mechanisms. Cell Biol Int 35:909–13.

34. Wiggins, C. H., D. Riveline, A. Ott, and R. E. Goldstein, 1998. Trapping and wiggling: elastohydrodynamics of driven microfilaments. Biophys J 74:1043–60.

35. Sader, J. E., 1998. Frequency response of cantilever beams immersed in viscous fluids with applications to the atomic force microscope. Journal of Applied Physics 84:64–76.

36. Lighthill, J., 1976. Flagellar Hydrodynamics - Neumann,Jv Lecture, 1975. Siam Review 18:161–230.

37. Svrcek-Seiler, W. A., I. C. Gebeshuber, F. Rattay, T. S. Biro, and H. Markum, 1998. Micromechanical Models for the Brownian Motion of Hair Cell Stereocilia. Journal of Theoretical Biology 193:623–630.

38. Weinbaum, S., X. Zhang, Y. Han, H. Vink, and S. C. Cowin, 2003. Mechanotransduction and flow across the endothelial glycocalyx. Proc Natl Acad Sci U S A 100:7988–95.

39. Guo, P., A. M. Weinstein, and S. Weinbaum, 2000. A hydrodynamic mechanosensory hypothesis for brush border microvilli. Am J Physiol Renal Physiol 279:F698–712.

40. Gao, Y., J. Wang, and H. Gao, 2010. Persistence Length of Microtubules Based on a Continuum Anisotropic Shell Model. Journal of Computational and Theoretical Nanoscience 7:1227–1237. https://doi.org/10.1166%2Fjctn.2010.1476.

41. Roth, K. E., C. L. Rieder, and S. S. Bowser, 1988. Flexible-substratum technique for viewing cells from the side: some in vivo properties of primary (9+0) cilia in cultured kidney epithelia. J Cell Sci 89 (Pt 4):457–66.

42. Mohieldin, A. M., W. A. AbouAlaiwi, M. Gao, and S. M. Nauli, 2015. Chemical-Free Technique to Study the Ultrastructure of Primary Cilium. Sci Rep 5:15982.

43. Rao, S. S., 2007. Vibration of continuous systems. Wiley, Hoboken, N.J.

44. Li, T., and M. G. Raizen, 2013. Brownian motion at short time scales. ANNALEN DER PHYSIK Annalen der Physik 525:281–295.

45. Tolic-Norrelykke, I. M., K. Berg-Sorensen, and H. Flyvbjerg, 2004. MatLab program for precision calibration of optical tweezers. Computer Physics Communications 159:225–240.

46. Happel, J., and H. Brenner, 1983. Low Reynolds number hydrodynamics: with special applications to particulate media. M. Nijhoff; Distributed by Kluwer Boston, The Hague; Boston Hingham, MA, USA.

47. Proudman, I., and J. R. A. Pearson, 1957. Expansions at small Reynolds numbers for the flow past a sphere and a circular cylinder. Journal of Fluid Mechanics 2:237–262.

48. Lamb, H., 1945. Hydrodynamics. Dover publications, New York„ 6th edition.

49. Van Dyke, M., 1975. Perturbation methods in fluid mechanics. Parabolic Press, Stanford, Calif., annotated edition.

50. Huang, T. C., 1964. Eigenvalues and modifying quotients of vibration of beams. University of Wisconsin, Engineering Experiment Station.

